# Plant-based production of SARS-CoV-2 antigens for use in a subunit vaccine

**DOI:** 10.1101/2021.10.17.464700

**Authors:** Jordan Demone, Maryam Nourimand, Mariam Maltseva, Mina Nasr-Sharif, Yannick Galipeau, Marc-André Langlois, Allyson M. MacLean

## Abstract

The COVID-19 pandemic has brought to the forefront an urgent need for the rapid development of highly efficacious vaccines, particularly in light of the ongoing emergence of multiple variants of concern. Plant-based recombinant protein platforms are emerging as cost-effective and highly scalable alternatives to conventional protein production. Viral glycoproteins, however, are historically challenging to produce in plants. Herein, we report the production of plant-expressed wild-type glycosylated SARS-CoV-2 Spike RBD (receptor-binding domain) protein that is recognized by anti-RBD antibodies and exhibits high-affinity binding to the SARS-CoV-2 receptor ACE2 (angiotensin-converting enzyme 2). Moreover, our plant-expressed RBD was readily detected by IgM, IgA, and IgG antibodies from naturally infected convalescent, vaccinated, or convalescent and vaccinated individuals. We further demonstrate that RBD binding to the ACE2 receptor was efficiently neutralized by antibodies from sera of SARS-CoV-2 convalescent and partially and fully vaccinated individuals. Collectively, these findings demonstrate that recombinant RBD produced *in planta* exhibits suitable biochemical and antigenic features for use in a subunit vaccine platform.

---

In this work, we describe the development of a rapid plant-based platform for the production of recombinant SARS-CoV-2 Spike Receptor-Binding Domain (RBD), which is simple, massively scalable, cost-effective and easily adaptable to reflect rapid changes in circulating viral sequences. Most importantly, our antigen closely recapitulates the biochemical and antigenic features of RBD produced in human cells, which is an essential feature for a plant-based human vaccine antigen. While most SARS-CoV-2 vaccines focus on intramuscular delivery systems, development of a subunit nasal spray vaccine offers a complementary method to increase mucosal immunity to SARS-CoV-2 variants and provide vaccine-hesitant and needle-phobic individuals with alternative immunization options. The RBD of the viral spike is the region primarily involved in binding to the cell surface receptor of the virus, the angiotensin converting enzyme 2 (ACE2) receptor (Hoffmann *et al*., 2020). Most neutralizing antibodies against SARS-CoV-2 are directed to the RBD (Garcia-Beltran *et al*., 2021; Piccoli *et al*., 2020; Yang *et al*., 2020).

A codon-optimized RBD sequence corresponding to the Wuhan-Hu-1 isolate of SARS-CoV-2 was cloned into the pHREAC vector backbone (Peyret *et al*., 2019) a plant-specific expression platform that has been demonstrated to achieve among the highest levels of *in vivo* protein expression reported in *N. benthamiana* (Fig. 1a). The RBD sequence was flanked on the N-terminal side by an endoplasmic reticulum targeting signal peptide derived from *Nicotiana tabacum* PR-1a signal peptide (amino acids 1-31), and on the C-terminal side by a tandem affinity tag consisting of an 8xHis tag and a Twin-Strep-tag (Fig. 1a). The dual affinity tags were separated by a glycine-serine linker, flanked by a thrombin cleavage site (LVPRGS) that was positioned directly upstream of the affinity tags, and an ER retention signal (KDEL) located downstream and adjacent to two tandem stop codons. Initial screening of the sequence validated expression vector demonstrated that the greatest expression was achieved using *Agrobacterium tumefaciens* strain AGL1 when infiltrated into five-to-six-week-old *N. benthamiana* leaves that were collected four days post-infiltration (dpi) (Fig. 1b). Margolin et al. 2020, had previously demonstrated much enhanced expression of a trimeric Spike mimetic protein in *N. benthamiana* when co-expressed in the presence of human calreticulin (Margolin, et al., 2020), thus we assessed expression of RBD in the presence and absence of this chaperone. Co-infiltrating with calreticulin (CRT) produced a noticeable increase in RBD expression levels (Fig. 1c) and we thus opted to co-express calreticulin with RBD in subsequent infiltrations with a consideration that the human chaperone may promote proper RBD folding. All RBD samples revealed a single band that migrated slightly above the expected molecular weight of 31.3 kDa, and slightly above a mammalian-expressed RBD control (Fig. 1d).

**Figure 1.**
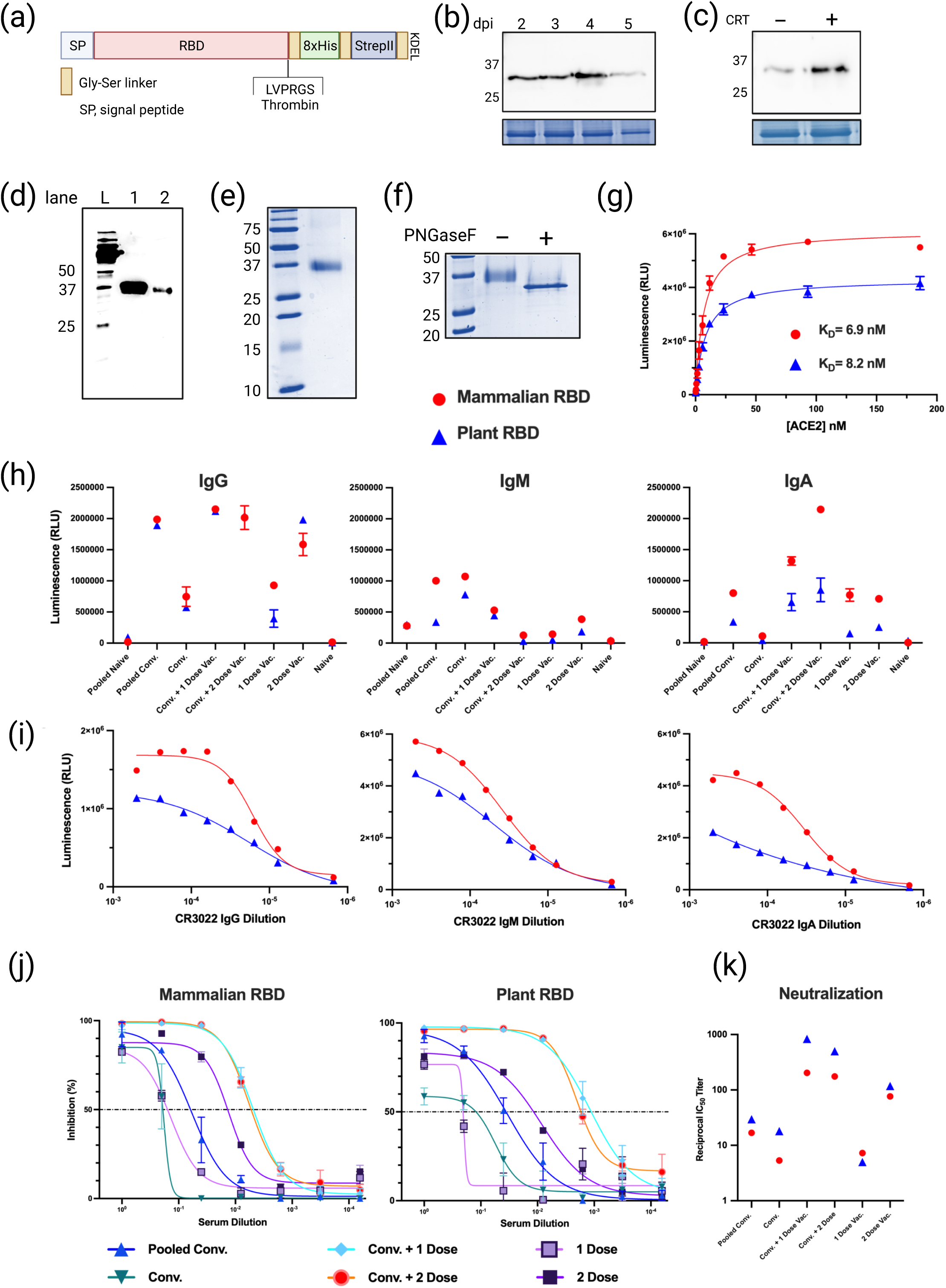
Purification and evaluation of SARS-CoV-2 Spike protein Receptor Binding Domain produced in *Nicotiana benthamiana*. (a) Schematic representation of genetic construct used to express SARS-CoV-2 RBD *in planta*. The SARS-CoV-2 sequence was expressed as a recombinant protein with a dual 8xHis and Twin-Strep II tag, interspersed with glycine-serine linkers (gold boxes). An ER-retention KDEL sequence was positioned at the C-terminus, and a Thrombin cleavage site (LVPRGS) was included for tag removal. SP, signal peptide. (b) Anti-His immunoblot of samples obtained from *N. benthamiana* 2 to 5 days post-infiltration (dpi) with RBD construct in (a). Loading control at 5 dpi reproducibly demonstrates reduced abundance of protein due to initiation of tissue necrosis at this time. (c) Co-infiltration of human calreticulin increases expression levels of RBD in *N. benthamiana*. Samples collected 4dpi. (d) Anti-his immunoblot of purified RBD expressed in *N. benthamiana* (lane 1) and control RBD expressed in mammalian 293F cells (lane 2). L, Bio-Rad protein ladder. (e) SDS-PAGE of RBD purified from *N. benthamiana*. (f) SDS-PAGE of plant-derived RBD treated with (+) and without (-) the amidase Peptide-N-Glycosidase F (PNGaseF), an enzyme that cleaves *N*-linked glycan chains. (g to k) A comprehensive assessment of protein function of the RBD produced in *N. benthamiana* as it pertains to protein folding and binding to ACE2 receptor, recognition and neutralization by antibodies in sera from SARS-CoV-2 exposed individuals. (g) Indirect ELISA demonstrating binding kinetics of soluble human ACE2 to immobilized mammalian RBD (red circle) and plant RBD (blue triangle). (h) Binding and recognition of immobilized mammalian and plant produced RBD by IgG, IgM and IgA polyclonal antibodies in sera of pooled naïve (unvaccinated, uninfected individuals; n>100), pooled convalescent (pooled Conv.) (n>100), convalescent and vaccinated with one dose (Conv. + 1 dose), convalescent and vaccinated with two doses (Conv. + 2 dose), vaccinated with one dose (1 dose), vaccinated with 2 doses (2 doses) of Pfizer (BNT162b2) and naïve (PCR negative confirmed). (i) Binding and recognition of immobilized mammalian and plant produced RBD by conformation dependent monoclonal IgG, IgM and IgA CR3022 antibodies. (j) Relative inhibition percentage of anti-SARS-CoV-2 neutralizing antibodies in blocking immobilized mammalian and plant produced RBD from binding soluble ACE2 by surrogate neutralization ELISA assay representative of technical triplicates or quadruplicates and presented as mean ± standard deviation. (k) Reciprocal half-maximal inhibitory dilution (ID_50_) values from (j) for mammalian and plant-produced RBD.

To scale up purification, we agroinfiltrated production batches of 25-30 five-to-six-week old plants (equivalent to approximately 200 grams fresh biomass) under a vacuum. Protein was partially purified by passing lysate through a Ni-NTA column, corresponding to purification of the 8xHis moiety of the tag. Fractions corresponding to the peak detected at 280 nm were visualized using immunoblot analysis to confirm presence and integrity of protein and were then loaded onto a StrepTrap column. Peak fractions eluting from that column were visualized via immunoblot, combined, concentrated, and digested overnight with thrombin to remove the 8xHis-Twin-Strep-tag. The digested sample was passed through the StrepTrap column a second time to remove the cleaved tag, and aliquots of the purified protein were analyzed on a Coomassie gel (Fig. 1e). A band on the Coomassie gel was visualized migrating at approximately 35 kDa, corresponding in migration to the band recognized on the RBD immunoblot by anti-His antibody. SARS-CoV-2 Spike protein has 22 *N*-linked glycosylation sites, two of which are located within RBD (Zhou *et al*., 2021). Peptide-*N*-glycosidase F (PNGase F) digestion of the partially purified protein confirmed that the plant produced RBD protein was glycosylated (Fig.1f).

We next performed a comprehensive assessment of the function and folding of RBD produced in *N. benthamiana*. Correct folding of the RBD antigen is critical for its interaction with the human ACE2 receptor and for the development of antibodies that will target and neutralize native virus and viral variants. To evaluate plant-produced RBD folding and binding kinetics, we first examined this interaction with the human ACE2 receptor. For this purpose, binding kinetics between soluble serially titrated ACE2 protein and immobilized plant RBD were evaluated by indirect enzyme-linked immunosorbent assay (ELISA). We found that plant-produced RBD readily binds to ACE2 with high affinity (K_D_=8.21 nM, R^2^=0.993), comparable to the RBD expressed in mammalian 293F cells (K_D_=6.90 nM, R^2^=0.981) (Fig. 1g).

Next, we examined whether plant-produced RBD was recognized by sera from COVID-19 convalescent, partially and fully vaccinated individuals. To this effect, sera from SARS-CoV-2 exposed individuals was added to immobilized plant RBD and detection by the three classes of antibodies induced by SARS-CoV-2 infection (IgG, IgM, and IgA) was investigated by ELISA. Here, we show that plant RBD was readily detected by IgM, IgA, and IgG antibodies from naturally infected convalescent, vaccinated or convalescent and vaccinated individuals (Fig. 1h). We reveal detection and high titers of IgG antibodies comparable to the mammalian produced RBD across all samples tested. We also report that IgM and IgA immunoreactivity to plant-produced RBD was generally less efficient in comparison to mammalian-derived RBD. Non-specific binding of antibodies to antigen-naïve individuals was minimal for both RBD antigens tested (Fig. 1h). Our plant-expressed RBD was recognized and bound by the conformation-dependent monoclonal IgM, IgA, and IgG CR3022 antibodies (Fig. 1i). This constitutes a further indirect indication that the plant RBD antigen is correctly folded and did not display significant inhibitory structural alterations or post-translation modifications. However, we did observe reduced binding of the CR3022 antibody to plant-derived RBD compared to mammalian-expressed RBD. These findings may suggest that post-translational modifications can impact antibody binding to some extent, which may also explain reduced binding of human IgM and IgA antibodies.

Lastly, we investigated the relative efficacy of human neutralizing antibodies in blocking the immobilized plant RBD’s interaction with soluble ACE2 by surrogate neutralization ELISA (Abe *et al*., 2020; Anderson *et al*., 2020). As mentioned previously, most neutralizing antibodies produced by natural exposure or vaccine immunity are directed to RBD. Evidence of neutralization is an additional indicator of adequate protein folding and antigenic epitope accessibility. Human serum was added to immobilized plant produced RBD and then binding to soluble biotinylated ACE2 receptor was evaluated (Fig. 1j). We then measured the inhibition of the binding as a function of the serum dilution. This enabled the calculation of the half-maximal inhibitory dilution (ID_50_) value. Our observations indicated that human antibodies were able to bind plant-expressed RBD and neutralize its binding to the ACE2 receptor with similar kinetics to the mammalian-expressed RBD (Fig. 1k). In fact, we observed generally greater ID_50_ values for the plant-expressed RBD, thereby indicating that its interactions with the ACE2 receptor were more easily disrupted by neutralizing antibodies. Given that affinity for ACE2 is very similar between human- and plant-produced RBD, greater ID_50_ values indirectly indicate that plant RBD exposes larger numbers of neutralizing epitopes, which could be due to differences in post-translational modifications such as glycosylation.

In summary, we have demonstrated that plant-produced RBD retains proper folding and functionality comparable to mammalian produced RBD as evaluated by binding to host cell receptor ACE2, recognition by the conformation-dependant monoclonal CR3022 antibody, and polyclonal antibodies from sera of SARS-CoV-2-infected individuals. Although we did note a reduced ability of IgM and IgA antibodies to bind plant-produced RBD as compared mammalian-expressed RBD, comparable binding of IgGs, which have high affinity to the target antigen and are the main antibody isotype involved in neutralization, were observed. Additionally, plant RBD binding to the ACE2 receptor was efficiently neutralized by antibodies from sera of SARS-CoV-2 convalescent and partially and fully vaccinated individuals. Collectively, these findings demonstrate that recombinant RBD produced in *N. benthamiana* exhibits suitable biochemical and antigenic features for use in a subunit vaccine platform.

## Acknowledgements

We are grateful for helpful discussions and advice from Rima Menassa and colleagues at Agriculture and Agri-Food Canada, London Research and Development Centre, Canada, Emmanuel Margolin, Edward Rybicki and colleagues from Biopharming Research Unit, University of Cape Town, South Africa, and Dominique Michaud, Université Laval, Canada. We gratefully acknowledge the excellent support of Michelle Brazeau, University of Ottawa, in her management of our greenhouse. MM holds a Queen Elizabeth II Graduate Scholarship in Science and Technology (QEII-GSST) and YG holds a Canadian Institute of Health Research (CIHR) Frederick Banting and Charles Best graduate scholarship (CGS-M). This study was supported in part by a COVID-19 Rapid Response grant by the Canadian Institute of Health Research (CIHR, #VR2 - 172722) and by a grant supplement by the Canadian Immunity Task Force (CITF) to MAL and funding from the University of Ottawa Faculty of Science to support greenhouse management. MAL holds a Canada Research Chair in Molecular Virology and Intrinsic Immunity.

## Author Contributions

AMM and MAL conceived and supervised the project. JD, MN, MNS optimized agroinfiltration and JD & MN optimized the purification protocol. JD performed the PNGaseF assay. MM and YG conducted *in vitro* functional assays and YG provided RBD control protein purified from HEK cells. JD, MN, MM, MAL and AMM contributed to the writing of the manuscript. All authors reviewed and approved the manuscript.

## METHODS

### Plant materials and growth conditions

Wild-type *Nicotiana benthamiana* plants were grown in a greenhouse with a 20h light/4h dark photoperiod at 25ºC for five weeks prior to infiltration. Plants were fertilized two to three times per week using Miracle-Gro (24-8-16) at a concentration of 1.1 g/L. Seeds were germinated on a weekly basis and 60 three-week-old seedlings were transplanted to individual pots once a week.

### Gene synthesis

The RBD sequence was obtained from the Wuhan strain (NC_045512). The sequence designed for expression in *N. benthamiana* had a C-terminal thrombin cleavage site, 8xhis tag, a Twin-Strep-tag®, and a KDEL sequence for ER-retention. The construct was codon-optimized for *N. benthamiana* expression and was synthesized by GenScript into the pHREAC vector (Peyret *et al*., 2019) via BsaI sites. Lectin-binding human chaperone calreticulin (NP_004334.1) was codon-optimized for *N. benthamiana* and synthesized in the pHRE vector (Peyret *et al*., 2019).

### Transient expression in *Nicotiana benthamiana*

Prior to infiltration, pHREAC-RBD and pHRE-calreticulin were freshly transformed into *Agrobacterium tumefaciens* strains AGL1 and Gv2260, respectively, via electroporation. A single colony was used to inoculate each 5 mL starter culture, and each culture was sub-cultured once in two litres of LB broth containing 50 mg/L kanamycin at 28ºC. The final cultures were grown to an optical density at 600nm (OD600) of 0.6 and were centrifuged and resuspended in MMA buffer (10 mM MES, 10 mM MgCl_2_, 200 μM acetosyringone, pH 5.6). *Agrobacterium* containing the pHREAC-RBD construct was resuspended to a final OD600 of 0.25 and the pHRE-construct was resuspended to a final OD600 of 0.1. The cultures were combined in a 1:1 volume ratio and incubated at room temperature for one hour prior to infiltration. Small-scale infiltration was performed by using 1 mL needleless syringes to inject *Agrobacterium* into the abaxial side of the leaves. Large-scale infiltration was performed via vacuum infiltration using three litres of infiltration solution. Leaves were gently scored, and individual plants were inverted and placed into the vacuum chamber for 2-3 minutes at a pressure of 5 bar. The vacuum was repeatedly applied until the plant was fully infiltrated. Infiltrated leaves were marked and plants were returned to the greenhouse until collection. Twenty-five plants were typically infiltrated per large-scale infiltration, equating to approximately 200 grams of fresh leaf biomass.

### Total soluble protein extraction

Leaves co-infiltrated with pHREAC-RBD and pHRE-calreticulin were collected 4-5 dpi (days post-infiltration). Leaf spines were removed, and tissue was homogenized in liquid nitrogen with mortar and pestle. Protein extracts were prepared under native conditions. For each FPLC run: 60 grams of ground tissue were resuspended in 270 mL of ice-cold buffer (PBS (20 mM NaH_2_PO_4_, 280 mM NaCl, 6 mM KCl), 10 mM imidazole, pH 7.4) and vortexed for five minutes. Ten microlitres of Lysonase Bioprocessing Reagent (Sigma Aldrich Canada Ltd; Etobicoke, Ontario) was added and the lysate was incubated at 4ºC for 15 minutes while gently shaking. Lysate were vortexed again and sonicated three times for 1 minute using a Kontes micro ultrasonic cell disruptor set at 70 output. Lysate was centrifuged for 45 minutes at 20,442 × *g* at 4ºC to pellet cell debris. Supernatants were filtered using several layers of cheesecloth and subsequently filtered using Corning one liter 0.22 μM polyethersulfone (PES) sterilizing low-binding filters (Corning, Incorporated; Corning, NY, USA).

### Purification of RBD using HisPur™ Ni-NTA Resin

Clarified total protein extract was passed through a HisPur Ni-NTA resin column (5 mL bed volume; Cytiva, Marlborough, US), pre-equilibrated with five volumes of imidazole binding buffer (PBS, 10 mM imidazole buffer, pH 7.4) using a fast protein liquid chromatography (FPLC) system (AKTA Pure; GE Healthcare systems). The column was washed with twenty bed volumes of binding buffer (PBS, 10 mM imidazole, pH 7.4) at a flow rate of 5 mL/minute. Protein was eluted using five bed volumes of elution buffer (PBS, 500 mM imidazole, pH 7.4) at a flow rate of 2 mL/minute. Samples were collected as 2 mL fractions. In initial experiments, aliquots of each fraction were visualized via immunoblotting (anti-His antibody; Cat: SAB2702218, Sigma Aldrich Canada, Oakville, Canada) to identify fractions enriched for RBD, which we found to consistently correspond to a visible peak eluting from the column in fractions 25 to 35.

### Purification of RBD using Strep-Tactin Resin

Fractions corresponding to the protein peak from the HisPur Ni-NTA resin were pooled and protein was further purified using a Strep-Tactin resin column (5 mL bed volume; Cytiva, Marlborough, US) using the AKTA Pure system. The Strep-Tactin column was pre-equilibrated with five bed volumes of binding buffer (PBS, pH 7.4) prior to sample loading, and the column was then washed with 10 bed volumes of binding buffer (PBS, pH 7.4). Protein was eluted using six bed volumes of StrepTrap elution buffer (PBS, 5 mM *d*-Desthiobiotin, pH 7.4) at a flow rate of 2 mL/minute. Samples were collected as 0.5 mL fractions. In initial experiments, aliquots of each fraction were immunoblotted (probed with anti-His antibody) and protein was also visualized via SDS-PAGE gels stained with Coomassie blue staining solution. We reproducibly detected RBD within fractions 22 to 33, which corresponded to the visible peak eluting from the column. Peak samples were collected, pooled, and concentrated to 500 μL using a Vivaspin 20 column (GE Healthcare; Mississauga, Canada).

### Further processing of RBD protein

Thrombin cleavage was performed to remove the tag from purified RBD protein. Ten microlitres of thrombin (1U / μL) were added to the 500 μL sample of purified RBD protein and incubated at 22ºC for 20 hours. To remove the cleaved 3 kDa tag from the final product, the reaction was passed through a Strep-Tactin resin column (5 mL bed volume) as described above. Flowthrough fractions corresponding to the peak were collected and concentrated to 500 μL using a Vivaspin 20 column (GE Healthcare, Mississauga, Canada). Protein concentration was quantified using the Bio-Rad Protein Assay and using BSA for standard curve preparation. Protein purity was analyzed using SDS-PAGE and the protein was visualized using Coomassie blue staining solution.

### PNGase F assay

PNGase F, an amidase, is an enzyme used to cleave *N*-linked glycans from glycoproteins and was used to assess the glycosylation status of plant-produced RBD. Briefly, 20 μg of partially purified RBD protein (following elution from Ni-NTA agarose) was incubated with denaturation buffer at 100ºC for 10 minutes. The sample was chilled to halt the denaturation process and GlycoBuffer 2 (New England Biolabs), NP-40, and PNGase F were added to the sample. The sample was incubated at 37ºC for one hour and analyzed on an SDS-PAGE gel. Bands were visualized using Coomassie blue staining solution.

### Indirect ELISA to assess ACE2 binding

To evaluate protein folding and binding, an indirect ELISA was carried out. Plant or mammalian produced SARS-CoV-2 RBD protein were diluted in sterile 1X PBS (Multicell #311-010-CL) to 4 μg/mL and coated onto a 384-well Immuno plates (Thermofisher, #460372) (12.5 μL /well) overnight at 4°C. Plates were washed three times with 100 μL of PBS-T (PBS, 1% Tween-20) using a BIOTEK plate washer (model ELX405) and blocked for one hour with blocking buffer (PBS-T + 3% non-fat milk powder, w/v) while shaking (700rpm) at room temperature. Biotinylated ACE2 as produced in Abe et al., 202013 was diluted in dilution buffer (PBS-T + 1% non-fat milk powder, w/v) to 258 ng/μL and subsequently 1:2 serially diluted. Plates were washed thrice with PBS-T, followed by addition of 20 μL per well of titrated soluble ACE2 and incubated for one hour while shaking at room temperature. Plates were washed thrice with PBS-T followed by the addition of 20 μL per well of Streptavidin-Peroxidase polymer (Sigma #S2438) diluted in dilution buffer to 1.25 ng/μL. After one hour incubation while shaking, plates were washed thrice with PBS-T and 20 μL of freshly prepared SuperSignal ELISA Pico Chemiluminescent Substrate (Thermo Scientific, #37069) (mixed 1:1 ratio and diluted in equal volume with dH_2_O, (V:V)) was added to each well. Following 5 mins of incubation on a shaker, luminescence signal (RLU) was measured with BIO-TEK Synergy Neo2 plate reader at 20 ms/well at a read height of 1.0 mm. Wells filled with dilution buffer in place of ACE2 accounted for background luminescence and were subtracted from the titrated ACE2 values. Dissociation constant (K_D_) was determined using 4-parameter curve fitting with GraphPad Prism 9.1.2 software.

### Indirect ELISA to evaluate anti-SARS-CoV-2 immunoreactivity in serum samples

Plant or mammalian produced SARS-CoV-2 RBD protein were diluted in sterile 1X PBS to 2 μg/mL and coated onto a 96-well plates (VWR #62402-959) (50 μL /well) overnight at 4°C. Plates were washed three times with 200 μL of PBS-T and blocked for one hour with a blocking buffer (PBS-T + 3% non-fat milk powder, w/v) on a shaker at room temperature. Serum samples were diluted 1:50 in dilution buffer (PBS-T + 1% non-fat milk powder, w/v). In conjunction, titration curves of conformation-dependent monoclonal IgM (Absolute Antibody, Ab01680-15.0), IgA (Absolute Antibody, Ab01680-16.0), and IgG (Absolute Antibody, Ab01680-10.0) CR3022 antibodies were used as reference material to assess protein folding. CR3022 antibodies were diluted 1:2000, followed by 1:2 serial dilution to establish a calibration curve. After blocking, plates were washed thrice with PBS-T, and followed by addition of 100 μL of the respective diluted serum samples and CR3022 antibodies. The plates were incubated for two hours on a shaker at room temperature, washed thrice with PBS-T followed by the addition of 50 μL of the respective secondary-HRP antibody at specified dilutions (1:4000 secondary anti-human IgG-HRP (NRC anti-hIgG#5-HRP fusion), anti-human 1:8000 IgA-HRP (Jackson ImmunoResearch, 109-035-011) or 1:9600 anti-human IgM-HRP (Jackson ImmunoResearch, 109-035-129)). Plates were incubated for one hour on a shaker, washed thrice with PBS-T followed by the addition with 100 μL of the diluted SuperSignal ELISA Pico Chemiluminescent Substrate. Luminescence intensity was measured with BIO-TEK Synergy Neo2 plate reader for 20ms/well at a read height of 1.0 mm. Wells filled with dilution buffer in place of serum accounted for background luminescence and were subtracted from the patient serum values.

### Surrogate Neutralization ELISA (snELISA) assay for evaluation of neutralization in serum samples

The described methodology was adapted from the surrogate neutralization ELISA assay as shown in Abe et al. 2020, for the evaluation of the relative inhibition of neutralizing antibodies to RBD protein from binding to soluble ACE2. Briefly, plant or mammalian produced SARS-CoV-2 RBD protein were diluted in sterile 1X PBS to 8 μg/mL and coated onto a 384-well Immuno plates (12.5 μL /well) overnight at 4°C. Plates were washed three times with PBS-T and blocked for one hour while shaking. Serum samples were 1:5 serially diluted in adilution buffer, applied to wells (20 μL/well) and incubated for 2 hours while shaking at room temperature. Plates were washed thrice, followed by the addition of biotinylated ACE2 diluted to 0.35 ng/μL in a dilution buffer (20 μL/well). Plates were washed thrice with PBS-T followed by the addition of 20 μL per well of Streptavidin-Peroxidase polymer diluted in dilution buffer to 1.25 ng/μL. Following one hour incubation, plates were washed thrice with PBS-T and freshly prepared SuperSignal ELISA Pico Chemiluminescent Substrate was applied (20 μL/well). Following 5 mins of incubation while shaking, luminescence intensity was measured with BIO-TEK Synergy Neo2 plate reader for 20ms/well at a read height of 1.0 mm. Control wells were filled with a dilution buffer in place of serum followed by the addition of ACE2 to assess maximum binding signal. Relative percent inhibition was calculated as follows:

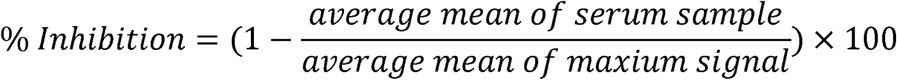

Serum dilution resulting in a 50% inhibition (half-maximal inhibitory dilution (ID_50_)) of RBD protein from binding ACE2 receptor was determined using 4-parameter fitting with GraphPad Prism 9.1.2 software.

### RBD Production in mammalian cells

For comparison, a mammalian RBD was produced in HEK 293F cells and purified. Briefly, a plasmid generously provided by Dr Florian Krammer (Mount Sinai, NYC) encoding the Whuhan-Hu-1 RBD (MN908947) sequence coding for the amino acid 319-541 and fused with the N-terminal SARS-CoV-2 spike secretory signal and a C-terminal hexa-histidine tag was transfected into 293F cells cultivated in Freestyle 293 expression media (Thermo Fisher, #12338018) at 37°C, 7% CO_2_, while shaking (125rpm). A total of 600 million cells resuspended in 200 mL were transfected with 200 μg of plasmid using ExpiFectamine (Thermo Fisher, 14525). Three days post-transfection, cell supernatant was harvested by centrifugation (4000xg for 20 mins at 4°C) and filtered through a low binding 0.22 μm stericup vacuum filter (Millipore Sigma, S2GPU10RE). The filtered supernatant was incubated for 2 hours at room temperature with 6ml of Ni-NTA resin (Qiagen, 30210). The column containing the mix of supernatant and resin was washed four times with a washing buffer containing 20mM imidazole, 300mM NaCl and 57.5mM of NaH_2_PO_4_·H_2_O. The RBD protein was then eluted with three column volumes of the elution buffer containing 234mM of imidazole, 300mM NaCl and 57.5mM of NaH_2_PO_4_·H_2_O. The eluted solution was concentrated and the buffer was replaced with PBS using a 10kDa Amicon filter (Millipore Sigma, UFC901008). RBD protein integrity was verified by SDS-PAGE, aliquoted to minimize freeze-thaw cycles and stored at -80°C.

### Patient Samples and Collection

Use of human samples for this study was approved by the University of Ottawa Ethics Review Board: Certificates H-04-20-5727, H-04-21-6643 and H-07-20-6009. The negative sample was taken from an individual with no history of SARS-CoV-2 infection and tested with PCR. Pooled negative samples were obtained from individuals negative for SARS-CoV-2 as tested by PCR and serology assay. Serum samples from convalescent or vaccinated patients enrolled in surveillance studies from different research studies post 2019. Samples were collected using standard phlebotomy procedures. Samples were de-identified and held at 4^0^C for short term handling and testing at University of Ottawa CL2+ biocontainment facility. All research was performed in accordance with current guidelines and regulations.

